# Speciation genomics and the role of depth in the divergence of rockfishes (*Sebastes*) revealed through Pool-seq analysis of enriched sequences

**DOI:** 10.1101/2022.06.06.494978

**Authors:** Daniel Olivares-Zambrano, Jacob Daane, John Hyde, Michael W. Sandel, Andres Aguilar

## Abstract

Speciation in the marine environment is challenged by the wide geographic distribution of many taxa and potential for high rates of gene flow through larval dispersal mechanisms. Depth has recently been proposed as a potential driver of ecological divergence in fishes and yet it is unclear how adaptation along these gradients’ shapes genomic divergence. The genus *Sebastes* contains numerous species pairs that are depth segregated and can provide a better understanding of the mode and tempo of genomic diversification. Here we present exome data on two species pairs of rockfishes that are depth segregated and have different degrees of divergence: *S. chlorostictus-S. rosenblatti* and *S. crocotulus-S. miniatus*. We were able to reliably identify ‘islands of divergence’ in the species pair with more recent divergence (*S. chlorostictus-S. rosenblatti*) and discovered a number of genes associated with neurosensory function, suggesting a role for this pathway in the early speciation process. We also reconstructed demographic histories of divergence and found the best supported model was isolation followed by asymmetric secondary contact for both species pairs. These results suggest past ecological/geographic isolation followed by asymmetric secondary contact of deep to shallow species. Our results provide another example of using rockfish as a model for studying speciation and support the role of depth as an important mechanism for diversification in the marine environment.

## Introduction

Understanding mechanisms of speciation in the marine environment remains difficult due to the lack of apparent geographical barriers and high rates of gene flow among populations (Dennenmoser et al. 2016). Most marine species demonstrate high dispersal capabilities and connectivity among populations, which can impede local adaptive processes and differentiation (Bierne et al. 2003, Carreras et al. 2017). However, even with the homogenizing effects of gene flow, there is potential for local adaptation that may be the driving force for genomic differentiation in the marine environment (Whitney et al. 2018).

Early work on speciation in marine fishes was thought to be a consequence of geographic isolating mechanisms. The formation of land barriers (Bermingham et al. 1997; Bernardi et al. 2004), islands (Leray et al. 2010), and physical boundaries generated from oceanographic processes (Hubert et al. 2012; Gaither and Rocha 2013) were used from a biogeographical perspective to describe speciation patterns in marine fishes. However, the role of pelagic larval duration in contributing to gene flow among populations suggested that allopatric divergence may be rarer in marine fish (reviewed in Bernardi 2013). Although large scale allopatric events can drive marine speciation, there is evidence that other isolating mechanisms occur in the marine environment (Rocha et al. 2005, Faria et al. 2020). Studies have demonstrated the possibility for closely related species to be sympatrically distributed, indicating ecology may be important (Rocha et al. 2005, Crow et al. 2010). Although Rocha et al. (2005) found evidence for sympatrically distributed reef fishes, there remains a lack of knowledge for how ecological speciation applies to temperate marine environments. Furthermore, sympatric or parapatric distribution of species contradicts existing knowledge that gene flow can be a barrier to speciation. This creates an apparent ‘marine speciation paradox’, or how can marine speciation occur in the face of high apparent gene flow (Johannesson 2009, Faria et al. 2020)?

The model of ecological speciation has become increasingly important in understanding the speciation process in the marine environment (Puebla 2009; Bernardi 2012). Numerous studies have now documented the role that ecological specialization, especially in fishes, plays in speciation. A number of environmental factors have been documented that lead to ecological divergence in the marine environment, these include temperature (Teske et al. 2019) and salinity (Momigliano et al. 2017), which are often correlated with other habitat characteristics (e.g. depth, latitude, etc.). Depth has already been identified as a potential factor in the diversification of rockfishes (Ingram 2011; Hyde et al. 2008; Heras and Aguilar 2019; Behrens et al. 2021), and depth may be important in driving speciation for other marine organisms (Carlon and Budd 2002; Prada and Hellberg 2013; Gaither et al. 2018; Hirase et al. 2021). Thus, adaptation to these environmental differences can lead to divergence in life history traits, such as spawning behavior, which subsequently drives reproductive isolation between incipient species.

Rockfishes (genus *Sebastes*) inhabit temperate waters across the Atlantic and Pacific Ocean, with sixty different species found in the North Pacific that have radiated over the past five million years (Johns and Avise 1998). Species are found from rocky intertidal habitats to depths greater than 1,500 m (Love et al. 2002). Given the ecological partitioning and habitat similarity between recently diverged forms of rockfish, ecological speciation may have contributed to their divergence (Behrens et al. 2021; Pavoine et al. 2009). Depth has been proposed as an important component in the diversification of rockfishes (Hyde et a. 2008; Ingram 2011.; Aguilar and Heras 2019; Behrens et al. 2021).

This study aims to identify genomic regions that have contributed to differentiation among recently diverged Northern Pacific species pairs of rockfish: (*S. chlorostictus-S. rosenblatti* and *S*.*-crocotulus-S*.*miniatus)*. The two species pairs occur along a continuum of divergence, with *S*.*-chlorostictus-S. rosenblatti* diverging approximately 0.21 Mya and *S*.*-crocotulus-S*.*miniatus* diverging approximately 2.3 Mya (Hyde and Vetter 2007). Both species pairs are found at different depths. *S*.*-chlorostictus* occurs between 60-240m while *S. rosenblatti* occurs between 100-490m. *S*.*miniatus* occurs at 30-100m while *S*.*-crocotulus* occurs between 100-200m (Hyde and Vetter 2007). Our goals are to determine if depth-segregated speciation is a result of selective sweeps that generates islands of genomic differentiation or ‘divergence islands’ (Via 2012, Wolf and Elegren 2016). We will also examine islands of genomic differentiation to see if they are shared across species pairs. Sharing of these divergence islands may indicate parallel evolutionary pressures in depth adaptation. Finally, we also investigated the demographic history of speciation in these species pairs, to see if similar patterns exist and if these patterns are consistent with ecological speciation.

## Methods and Analysis

### Sample Collection

Ethanol preserved fin clips for the following depth-separated species pairs were obtained from *S chlorostictus-S. rosenblatti* and *S. crocotulus-S. miniatus* (Table 1). High molecular weight DNA was obtained using standard phenol-chloroform methods followed by ethanol precipitation (Sambrook and Russel 2006). DNA integrity was checked on a 2% agarose gel and quantified on a Qubit fluorometer before preparing the samples for Illumina library preparation.

**Table 1:**
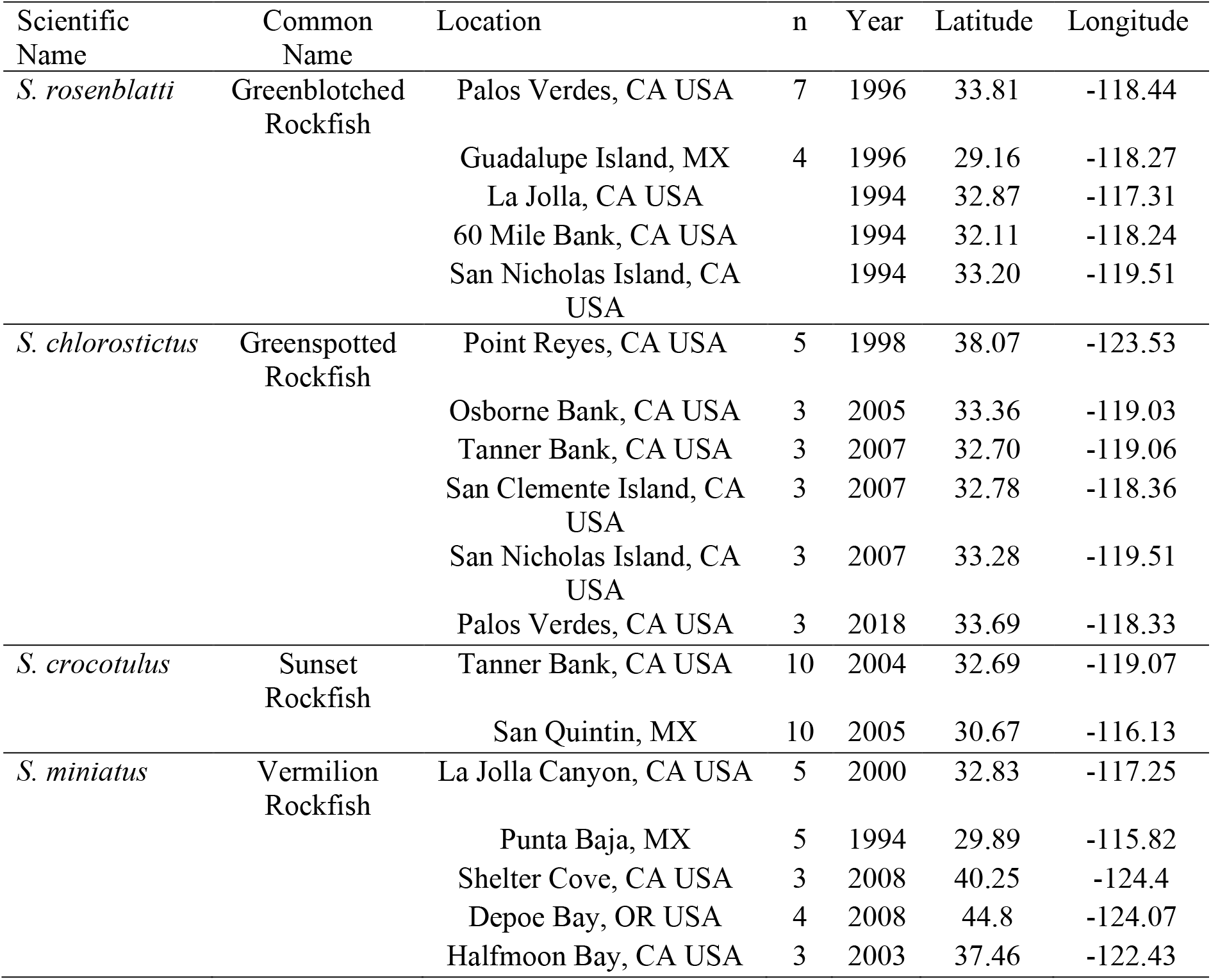
Rockfish samples used for this study.

| Scientific Name | Common Name | Location | n | Year | Latitude | Longitude |
| --- | --- | --- | --- | --- | --- | --- |
| *S. rosenblatti* | Greenblotched Rockfish | Palos Verdes, CA USA | 7 | 1996 | 33.81 | -118.44 |
|  |  | Guadalupe Island, MX | 4 | 1996 | 29.16 | -118.27 |
|  |  | La Jolla, CA USA |  | 1994 | 32.87 | -117.31 |
|  |  | 60 Mile Bank, CA USA |  | 1994 | 32.11 | -118.24 |
|  |  | San Nicholas Island, CA USA |  | 1994 | 33.20 | -119.51 |
| *S. chlorostictus* | Greenspotted Rockfish | Point Reyes, CA USA | 5 | 1998 | 38.07 | -123.53 |
|  |  | Osborne Bank, CA USA | 3 | 2005 | 33.36 | -119.03 |
|  |  | Tanner Bank, CA USA | 3 | 2007 | 32.70 | -119.06 |
|  |  | San Clemente Island, CA USA | 3 | 2007 | 32.78 | -118.36 |
|  |  | San Nicholas Island, CA USA | 3 | 2007 | 33.28 | -119.51 |
|  |  | Palos Verdes, CA USA | 3 | 2018 | 33.69 | -118.33 |
| *S. crocotulus* | Sunset Rockfish | Tanner Bank, CA USA | 10 | 2004 | 32.69 | -119.07 |
|  |  | San Quintin, MX | 10 | 2005 | 30.67 | -116.13 |
| *S. miniatus* | Vermilion Rockfish | La Jolla Canyon, CA USA | 5 | 2000 | 32.83 | -117.25 |
|  |  | Punta Baja, MX | 5 | 1994 | 29.89 | -115.82 |
|  |  | Shelter Cove, CA USA | 3 | 2008 | 40.25 | -124.4 |
|  |  | Depoe Bay, OR USA | 4 | 2008 | 44.8 | -124.07 |
|  |  | Halfmoon Bay, CA USA | 3 | 2003 | 37.46 | -122.43 |

### Targeted Sequence Capture Design

We designed a series of oligonucleotide capture baits that could efficiently enrich DNA sequencing libraries across Perciformes, a large order of more than 2,200 species that includes *Sebastes* (Daane et al. 2016; Nelson et al. 2016; Daane et al. 2019). We targeted protein coding exons, conserved non-coding elements (CNEs), miRNAs and ultra-conservative elements (UCNEs) for enrichment. Protein coding exons were extracted from Ensembl BioMart for the three-spined stickleback (*Gasterosteus aculeatus*, BROAD S1), the Japanese medaka (*Oryzias latipes*, MEDAKA1) and green spotted puffer (*Tetraodon nigroviridis*, TETRAODON 8.0 (Kinsella et al. 2011). CNEs were defined from the constrained regions > 50bp within the Ensembl compara 11-way teleost alignment that did not overlap with coding sequences (Ensembl release-91)(Herrero et al. 2016). miRNA hairpins were identified from miRbase and UCNEs from UCNEbase (Kozomara et al. 2011; Dimitrieva et al. 2013).

We used BLASTN (ncbi-blast-2.2.30+; parameters ‘-max_target_seqs 1-outfmt 6’) to identify each targeted element within multiple perciform genome assemblies. The majority of capture baits were designed against the genome of the Chabot de Rhénanie *Cottus rhenanus* (ASM145555v1). Importantly, certain genetic regions may be absent or highly divergent within the genome of this sculpin but remain conserved in other Perciformes. To account for these regions and ensure their capture, we iteratively designed capture baits from the genomes of the shorthorn sculpin *Myoxocephalus scorpius* (ASM90031295v1) (Malmstrøm et al. 2016), the sablefish *Anoplopoma fimbria* (AnoFim1.0)(7), the golden redfish *S. norvegicus* (ASM90030265v1) (Malmstrøm et al. 2016), the flag rockfish *S. rubrivinctus* (SRub1.0), the rougheye rockfish *S. aleutianus* (ASM191080v2), the European perch *Perca fluviatilis* (ASM90030264v1) (Rondeau et al. 2013), and the three-spined stickleback *G. aculeatus* (BROAD S1). For each species, we included new capture baits if the targeted elements were either not identified (coverage < 70% or a BLASTN E-value >0.001), or had < 85% identity to an existing capture bait. As a result of this iterative addition of sequence information from new species, there should be oligonucleotide capture baits of at least 85% identity to each clade included in the capture design. This multi-species design enables efficient enrichment across distantly-related perciform fishes. The final species composition of the capture baits: *C. rheanus* (62.0%), *M scorpius* (6.7%), *A. fimbria* (6.4%), *S. norvegicus* (5.9%), *S. rubrivinctus* (2.0%), *S. aleutianus* (1.8%), *P. fluviatilis* (5.3%), *G. aculeatus* (9.8%)

SeqCap EZ Developer (cat #06471684001) capture oligos were designed in collaboration with the Nimblegen design team to standardize oligo annealing temperature, reduce probe redundancy, and to remove low complexity DNA regions. The capture design contained sequence from 492,506 regions (81,493,221 total bp) across all 8 perciform reference genomes. Accounting for probe redundancy between the perciform reference genomes, the final capture design comprised 407,084 distinct elements, including 285,872 protein coding exons, 118,406 conserved non-coding elements, 2,508 UCNEs and 298 miRNAs (see Daane et al. 2016).

### Exome Sequencing

Exome sequencing of 20 individuals from four different species was done using a pool-seq approach (Table 1) (Schlotterrer et al. 2014; Daane et al 2016). DNA from 20 individuals was pooled in equimolar amounts and libraries were constructed using the Kapa HyperPlus kit (Supplemental Table S1). Enrichment of exome sequences was done following the approach of Daane et al. (2019). Individually barcoded libraries were quantified using qPCR, pooled, and sequenced on an Illumina HiSeq4000 at the UC Berkeley Vincent Coates Genomics facility with 150 PE sequencing.

### Read Mapping

Following sequencing, reads were demultiplexed, then trimmed and quality filtered with Trimmomatic (Bolger et al. 2014) using the parameters: ILLUMINACLIP:TruSeq3-PE-2:2:30:10 LEADING:3 TRAILING:3 SLIDINGWINDOW:4:15 MINLEN:36” ILLUMINACLIP:TruSeq3-PE-2:2:30:10 LEADING:3 TRAILING:3 SLIDINGWINDOW:4:15 MINLEN:36. The resulting high-quality trimmed reads were mapped to the *S. umbrosus* genome (Kolora et al. 2021; assembly fSebUmb1.pri) with bwa-mem (Li and Durbin 2010). Resulting BAM files were converted to mpileup format with samtools (Supplemental Table S1) (Danecek et al. 2021) and regions surrounding indels were masked with the identify-indel-regions.pl script for subsequent analysis. Allele frequencies were estimated with Popoolation2 (Kofler et al. 2011) and loci with a minor allele frequency (MAF) less than 0.05 were removed for subsequent analyses.

We identified divergence islands across species pairs using outlier approaches. We estimated F_ST_ for species pairs using the parameters in Popoolation2: --suppress-noninformative --min-count 6 --min-coverage 80 --max-coverage 500 --min-covered-fraction 1 --window-size 1 --step-size 1 --pool-size 20 (Li et al. 2009, Kofler et al. 2011). Results from the F_ST_ analysis were then plotted utilizing qqman to generate Manhattan Plots in R (Turner 2018).

### Sliding Window Analysis

To identify divergence islands we applied an approach similar to Renaut et al. (2013) and Holliday et al. (2016). We performed a sliding window analysis of F_ST_ across each chromosome, using a window of 10 adjacent SNPs and sliding the window every two SNPs, to identify the number of regions that contained SNPs greater than the top one percentile for F_ST_ of the genome-wide analysis. To assess significance, we randomly sampled 10 SNPs from across the genome with replacement for their F_ST_ values 100,000 times. For each of these subsamples we estimated the proportion of top one percentile F_ST_ SNPs present. The proportion of top one percentile SNPs in each window, over the resampled dataset, was used to determine significance for the original dataset (p <0.001).

### Gene Ontology Enrichment Analysis

We identified genes found in outlier windows from our sliding window analysis. These served as our candidate list of genes that was then compared to the background list of all annotated genes found from our exome dataset. To test for enrichment of molecular pathways and/or function between the background and candidate lists and to allow for the visualization of the gene ontology networks we used Cytoscape-CLUEGO and its plug in CLUEPedia (Bindea et al. 2009, Bindea et al. 2013). Cytoscape-CLUEGO utilizes hypergeometric testing followed by Bonferroni multiple testing corrections between a candidate gene list and a custom background list (Bindea et al. 2009). For Cytoscape-CLUEGO, all annotations were made using the *D. rerio* genome from the Gene Ontology Consortium (Ashburner et al. 2000), provided as it was the closest related species to genus *Sebastes* in this analysis package. Finally, Cytoscape-CLUEGO groups genes by GOterm to avoid redundancy in the results.

### Demography of Speciation

We utilized folded site frequency spectra in δaδi (Gutenkunst et al. 2009) to determine the demographic history of speciation in each of the species pairs. To generate datasets for this analysis we used all identified SNPs from Popoolation2 (before filtering for MAF as above). To assure independence of each SNP, we randomly sampled one SNP every 1,000,000 bp across the genome. Raw SNP frequencies were converted into δaδi SNP format using genomalicious (https://rdrr.io/github/j-a-thia/genomalicious/). We explored seven simple two-population models in δaδi: no migration, symmetric migration, asymmetric migration, symmetric migration followed by isolation, asymmetric migration followed by no migration, secondary contact with symmetric migration, and secondary contact with asymmetric migration. We hypothesize that if ecological speciation has occurred in these species pairs, due to adaptation to different depths, we would expect to observe a demographic history with at least some gene flow. We utilized the δaδi optimization procedure from Portik et al. (2017) (https://github.com/dportik/dadi_pipeline). We ran four iterations of optimizations for each model with 10, 20, 30, and 40 replications respectively. For each species pair we compared models using AIC and Δ AIC. We did not convert parameters from the best fit model into biologically meaningful values, as our goal was simply to reconstruct a reasonable demographic scenario for each species pair.

## Results

### Genome Assembly and Coverage

We obtained sequences for pooled sequenced from each species that contained 20 individuals. The average read depth was 10.36 across species and > 99% of the reads mapped back to the *S. umbrinus* genome (Supplemental Table S1).

### SNP Calling and F_ST_

We used F_ST_ to identify islands of divergence between each species pair and found F_ST_ values for 48,106 SNPs in *S. crocotulus-S. miniatus* and 52,626 SNPs in *S. chlorostictus-S. rosenblatti*. Mean F_ST_ for *S. crocotulus-S. miniatus* was 0.10 with a standard error of 0.0014; for *S. chlorostictus-S. rosenblatti*, mean F_ST_ was 0.03 with a standard error of 0.0003. We found a total of 10 non-overlapping windows in the *S. crocotulus-S. miniatus* comparison that passed our significance threshold (p < 0.0001) and 33 windows for *S. chlorostictus-S. rosenblatti* (Figure 1; Supplemental Tables S2 and S3). There were two windows that were shared in both comparisons, one on chromosome 6 and the other on chromosome 12 (Figure 1, Supplemental Table S2-S4).

### Enrichment Analysis of Significant F_ST_ Windows and Candidate Genes

Genes found within significant F_ST_ windows for *S*.*chlorostictus-S. rosenblatti* were enriched for pathways related to neuropeptide signaling, monovalent inorganic cation inorganic homeostasis, galanin receptor activity, proton-transporting two-sector ATPase complex - catalytic domain, active transmembrane transporter activity, P-P-bond-hydrolysis driven transmembrane transporter activity, and proton transmembrane transporter activity (Bonferroni < 0.05). Eighteen of the 200 candidate genes in this species pair were enriched within these GO terms. Of the 18 genes: five are related to ATP binding, ATP synthase or ATPase activity (*abcb8, atp5f1e, atp6v1h, atp1b2b, atp6ap1a*), four are solute carriers (*slc4a2b, slc12a9, slc2a10, slc9a7*), two are integrin beta subunits (*itb4r, itb4r2a*). The remaining seven genes have functions related to melanophore production stimulation, germ cell migration, behavioral, ectodermal placode development, cell proliferation inhibition, and MHC class I binding activity (*adcyap1b, ca15b, galr1b, oprk1, pnocb, pth2, tap2t*) (Table 2, Figure 2A, Supplementary Table S5).

**Table 2:**
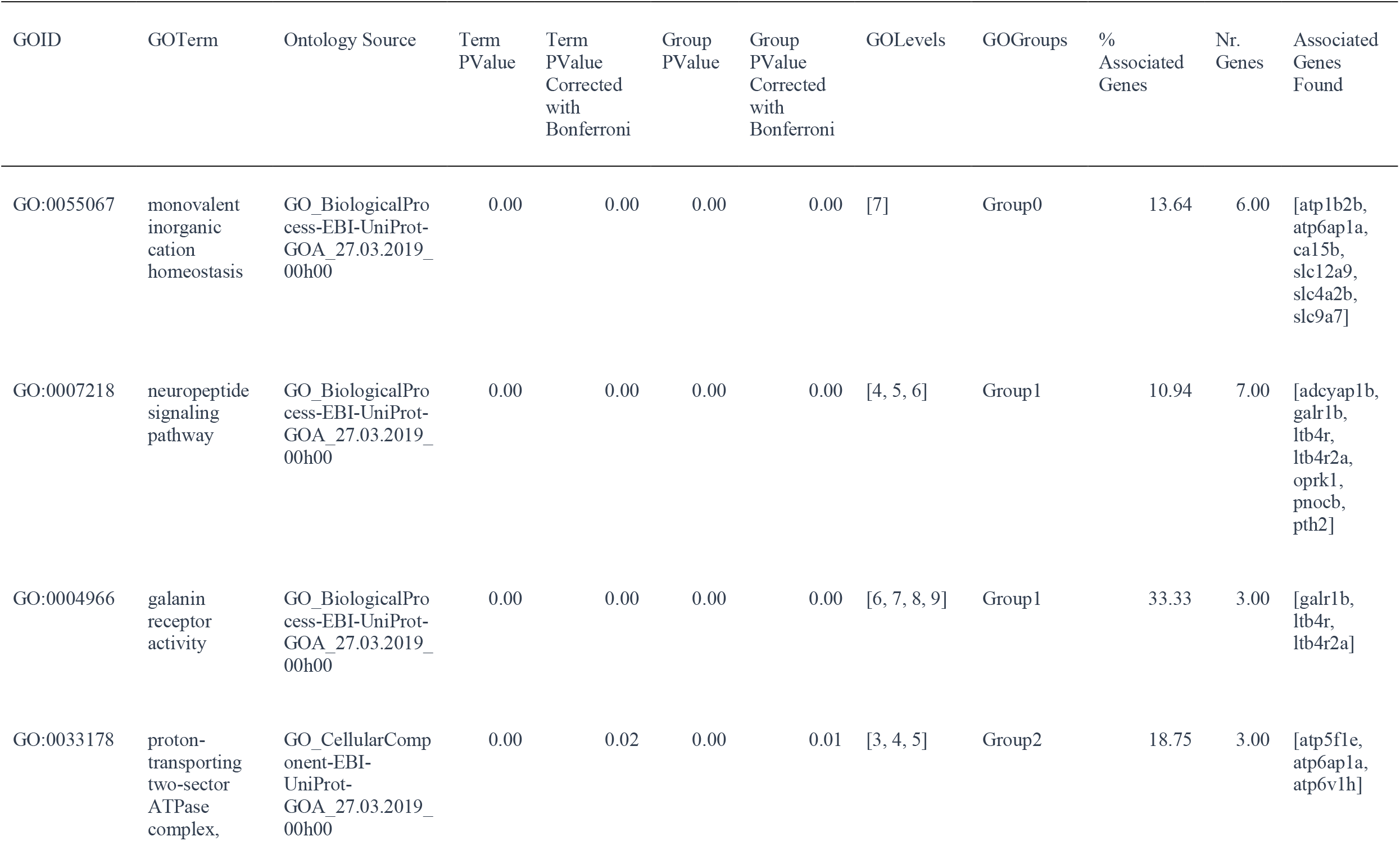
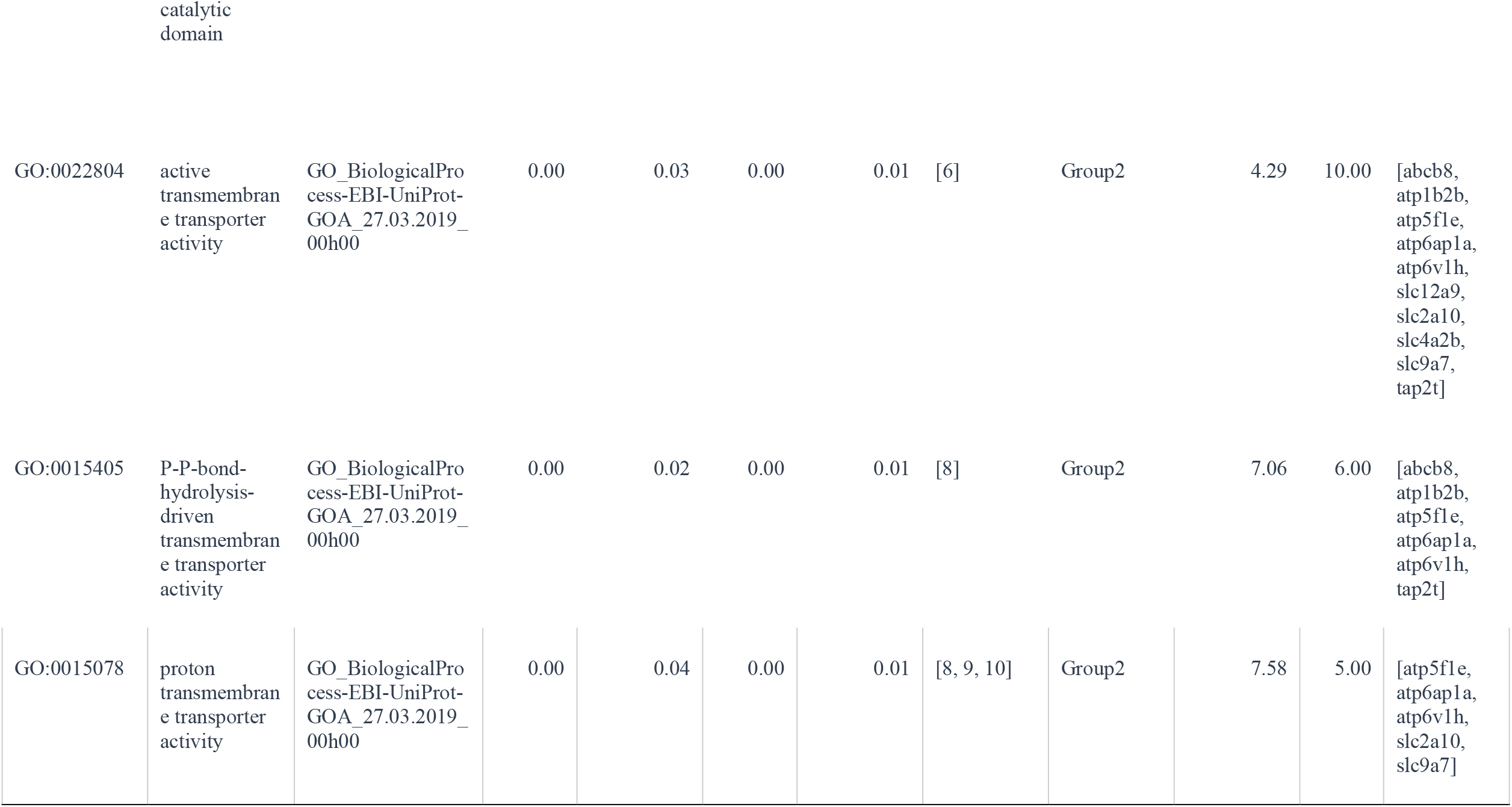
Enriched GO Terms in *S. chlorostictus-S. rosenblatti* candidate genes found near genomic islands of divergence utilizing Cytoscape-CLUEGO (Bonferroni < 0.05).

**Figure 1.**
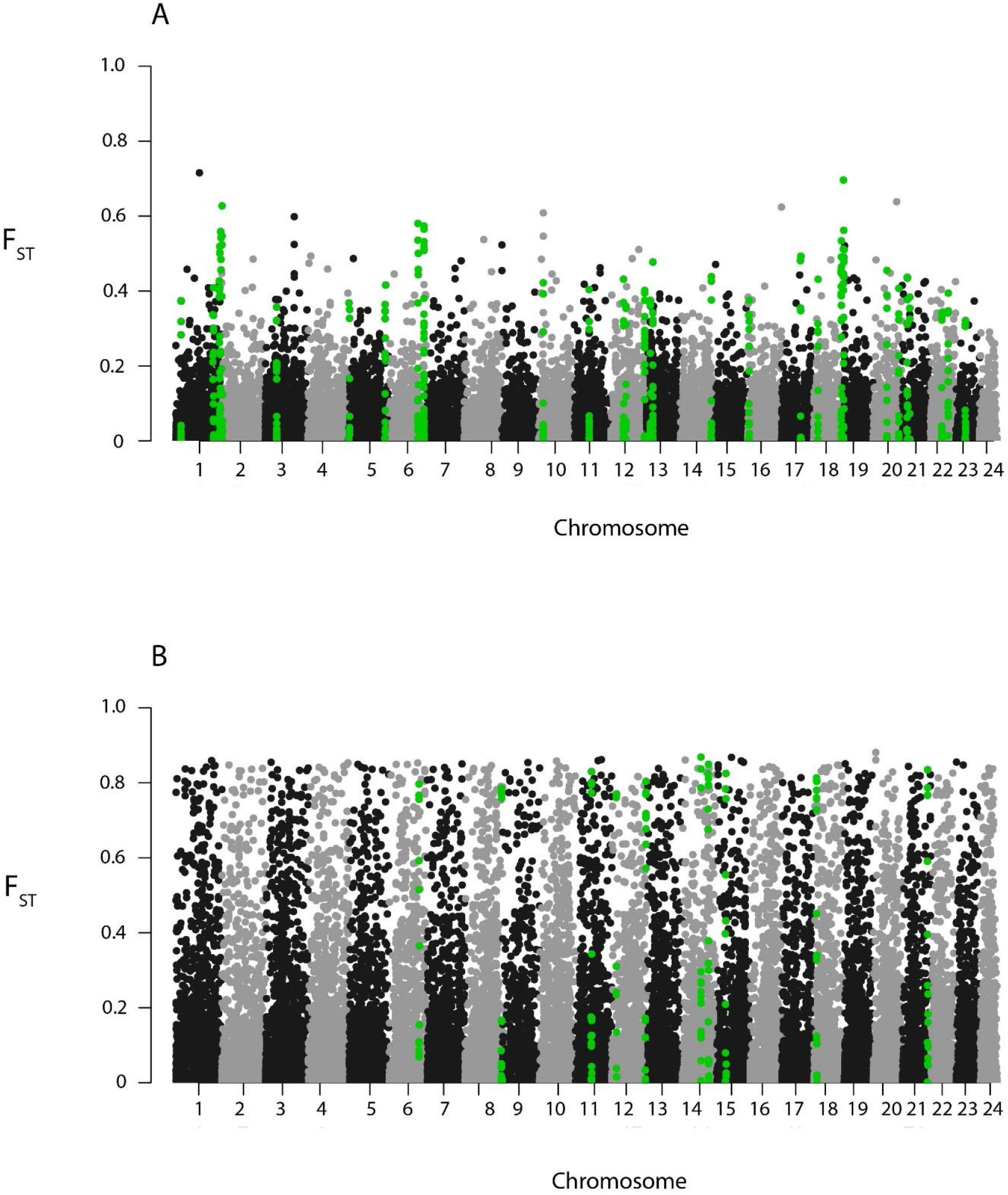
Manhattan plot for *S. chlorostictus-S. rosenblatti* (A) and *S. crocotulus-S. miniatus* (B) species pairs displaying individual SNP F_ST_ values by chromosome number (aligned to S. umbrosus genome). Highlighted green values are significant windows based on analyses described in the methods.

**Figure 2:**
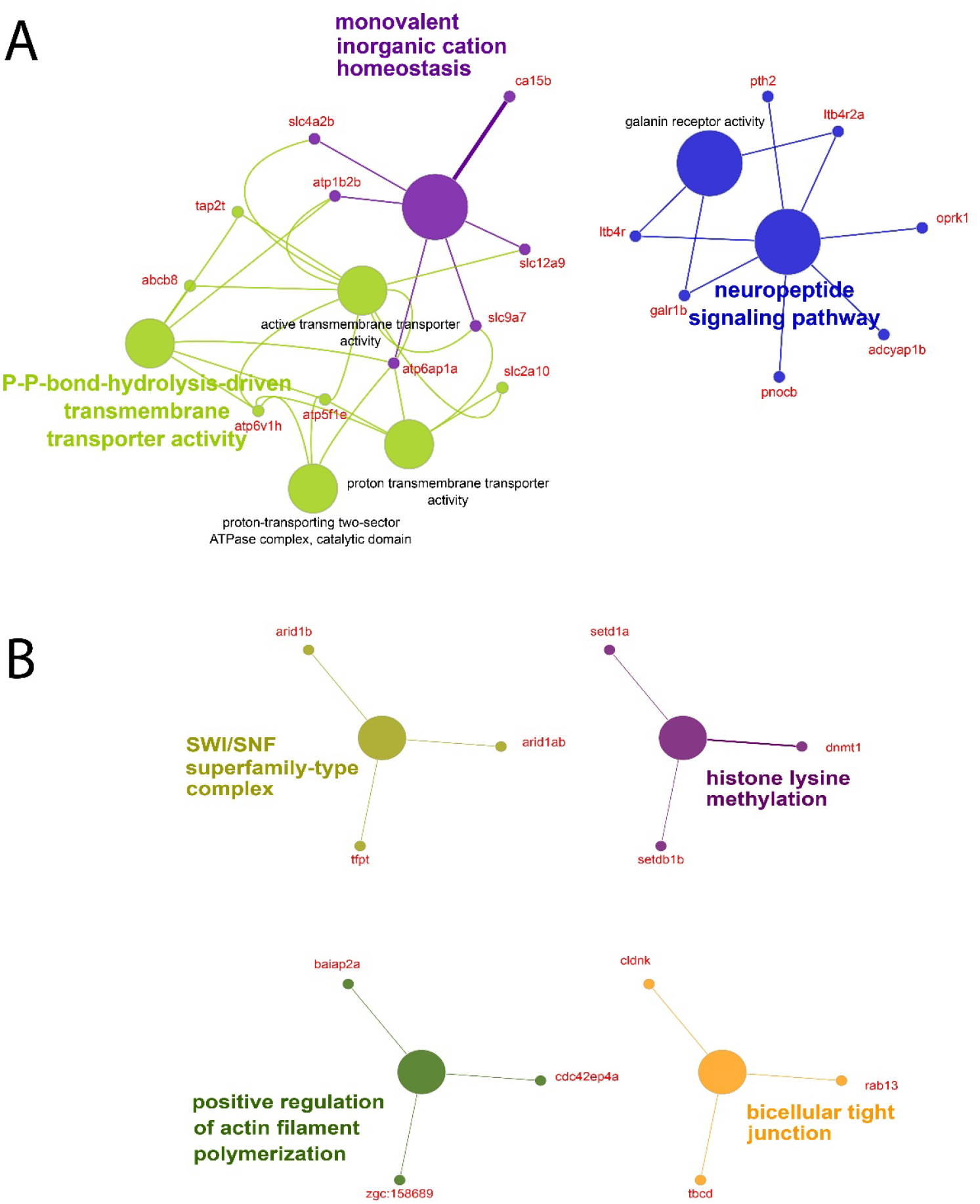
Significant GO terms for genes in genomic islands of divergence identified via sliding window analysis (Bonferonni < 0.05) for the *S. chlorostictus-S. rosenblatti* (A) and *S. crocotulus-S. miniatus* (B) comparisons. The different colors shaded in the circles represent unique functional GO terms. Linked functional GO terms illustrate a functional pathway. The candidate genes that connect each significant GO term are labeled red. Finally, highlighted terms are known as a leading term as they are the most significant GO term from the analysis.

Genes found within significant F_ST_ windows for *S*.*crocotulus-S. miniatus* were enriched for pathways related to bicellular tight junction, positive regulation of actin filament polymerization, histone lysine methylation and SWI/SNF superfamily-type complex (Bonferroni < 0.05). Alone, 12 candidate genes from a list of 126 candidate genes were enriched for these GO terms and provide functional and cellular pathway insight. Of the 12, three are involved in chromatin remodeling (*arid1ab, arid1b, tfpt*), three are involved in methylation (*dnmt1, setd1a, setdb1b*), four are related to cytoskeletal organization (*zgc:158689, tbcd, cldnk, baiap2a*), one to GTP binding (*cdc42ep4a*) and one to intracellular membrane organization (*rab13*) (Table 3, Figure 2B, Supplementary Table S5).

**Table 3:**
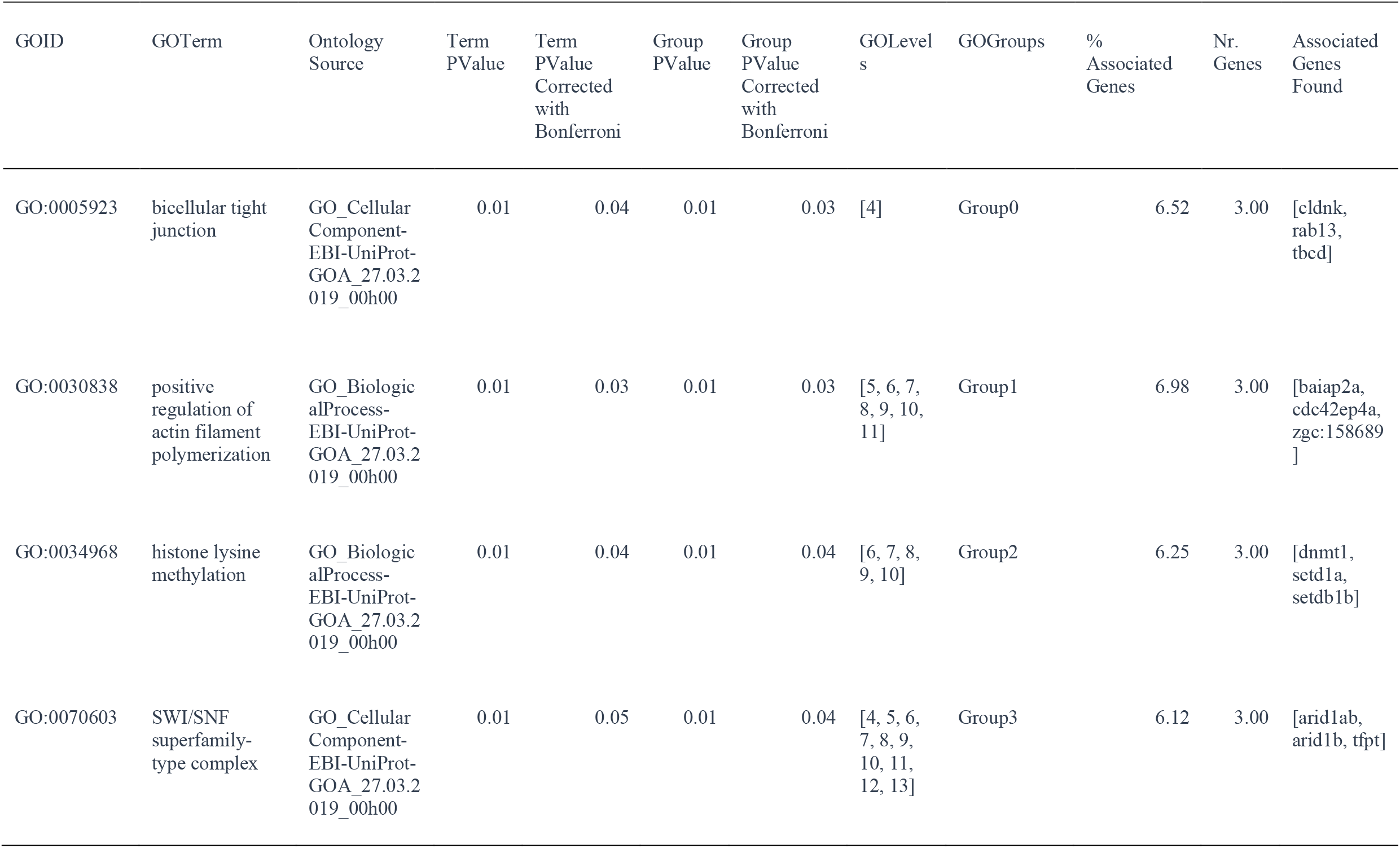
Enriched GO Terms in *S. crocotulus-S. miniatus* candidate genes found near genomic islands of divergence utilizing Cytoscape-CLUEGO (Bonferroni < 0.05).

| GOID | GO Term | Ontology Source | Term PValue | Term PValue Corrected with Bonferroni | Group PValue | Group PValue Corrected with Bonferroni | GO Levels | GO Groups | % Associated Genes | Nr. Genes | Associated Genes Found |
| --- | --- | --- | --- | --- | --- | --- | --- | --- | --- | --- | --- |
| GO:0005923 | bicellular tight junction | GO_Cellular Component-EBI-UniProt-GOA_27.03.2019_00h00 | 0.01 | 0.04 | 0.01 | 0.03 | [4] | Group0 | 6.52 | 3.00 | [cldnk, rab13, tbcd] |
| GO:0030838 | positive regulation of actin filament polymerization | GO_Biological Process-EBI-UniProt-GOA_27.03.2019_00h00 | 0.01 | 0.03 | 0.01 | 0.03 | [5, 6, 7, 8, 9, 10, 11] | Group1 | 6.98 | 3.00 | [baiap2a, cdc42ep4a, zgc:158689] |
| GO:0034968 | histone lysine methylation | GO_Biological Process-EBI-UniProt-GOA_27.03.2019_00h00 | 0.01 | 0.04 | 0.01 | 0.04 | [6, 7, 8, 9, 10] | Group2 | 6.25 | 3.00 | [dnmt1, setd1a, setdb1b] |
| GO:0070603 | SWI/SNF superfamily-type complex | GO_Cellular Component-EBI-UniProt-GOA_27.03.2019_00h00 | 0.01 | 0.05 | 0.01 | 0.04 | [4, 5, 6, 7, 8, 9, 10, 11, 12, 13] | Group3 | 6.12 | 3.00 | [arid1ab, arid1b, tfpt] |

### Demography of Speciation

In order to assess the demographic history of speciation within our two species pairs we tested seven models of population divergence using δaδi. We used pruned datasets that consisted of 5,368 and 5,310 SNPS for the *S. rosenbaltti-S. chlorostictus* and *S. miniatus-S. crocotulus* species pairs respectively. For both species pairs we found the secondary contact with asymmetric geneflow from the deep to shallow species to be the best model (Tables 4 and 5).

**Table 4:** Results of the demographic model analysis (δaδi) for the *S. chlorostictus-S. rosenblatti* species pair

| Model abbreviation | k | Log-l | AIC | delta AIC | chi-squared | theta | nu1 | nu2 | m | m1->2* | m2->1* | T | T1 | T2 |
| --- | --- | --- | --- | --- | --- | --- | --- | --- | --- | --- | --- | --- | --- | --- |
| *sec_contact_asym_mig* |  | -2558.41 | 5128.82 | 0 | 5653.84 | 997.87 | 0.6277 | 0.1753 |  | 0.1814 | 9.157 |  | 7.5564 | 0.3685 |
| *anc_asym_mig* |  | -2654.33 | 5320.66 | 191.84 | 4537.62 | 538.23 | 1.3647 | 0.7972 |  | 0.1427 | 1.7962 |  | 3.1908 | 0.0143 |
| *no_mig* |  | -2673.41 | 5352.82 | 224 | 4641.79 | 1366.47 | 0.1264 | 0.1427 |  |  |  | 0.0275 |  |  |
| *asym_mig* |  | -2676.33 | 5362.66 | 233.84 | 4642.29 | 782.12 | 0.9455 | 0.5535 |  | 0.1723 | 2.3527 | 0.9105 |  |  |
| *sym_mig* |  | -2780.87 | 5569.74 | 440.92 | 4583.46 | 215.43 | 2.6643 | 3.0234 | 0.261 |  |  | 5.7288 |  |  |
| *sec_contact_sym_mig* |  | -2780.65 | 5571.3 | 442.48 | 4591.22 | 173.31 | 3.3009 | 3.7447 | 0.2146 |  |  | 6.4535 |  |  |
| *anc_sym_mig* |  | -2845.23 | 5700.46 | 571.64 | 5855.37 | 335.37 | 1.7422 | 1.7978 | 2.296 |  |  |  | 3.7806 | 0.2338 |

\* - 1: *S. rosenblatti*; 2: *S. chlorostictus*

**Table 5:** Results of the demographic model analysis (δaδi) for the S. crocotulus-S. minatus species pair

| Model abbreviation | k | Log-l | AIC | delta AIC | chi-squared | theta | nu1 | nu2 | m | m1->2* | m2->1* | T | T1 | T2 |
| --- | --- | --- | --- | --- | --- | --- | --- | --- | --- | --- | --- | --- | --- | --- |
| *sec_contact_asym_mig* |  | -1799.93 | 3611.86 | 0 | 2443.07 | 151.38 | 0.6573 | 2.5635 |  | 9.0238 | 1.0233 |  | 28.6354 | 0.4318 |
| *no_mig* |  | -1895.39 | 3796.78 | 184.92 | 2105.57 | 1246.52 | 0.0311 | 0.0544 |  |  |  | 0.0022 |  |  |
| *sym_mig* |  | -2083.7 | 4175.4 | 563.54 | 2375.96 | 114.23 | 4.5392 | 5.5264 | 0.8282 |  |  | 20.2339 |  |  |
| *asym_mig* |  | -2082.8 | 4175.6 | 563.74 | 2400.21 | 260.87 | 1.6168 | 2.8042 |  | 2.5446 | 1.4564 | 8.119 |  |  |
| *sec_contact_sym_mig* |  | -2087.1 | 4184.2 | 572.34 | 2407.4 | 325.39 | 1.6042 | 1.8633 |  | 2.5214 |  |  | 2.9266 | 2.7467 |
| *anc_asym_mig* |  | -2112 | 4236 | 624.14 | 2539.94 | 145.73 | 2.2964 | 6.0197 |  | 2.5924 | 0.6727 |  | 17.1097 | 0.0377 |
| *anc_sym_mig* |  | -2230.2 | 4470.4 | 858.54 | 2884.82 | 1071.12 | 0.3464 | 0.8302 | 0.6722 |  |  |  | 0.0132 | 0.0119 |

\* - 1: *S. crocotulus*; 2: *S. miniatus*

## Discussion

The presumed lack of geographic barriers and propensity for high levels of gene flow in poses challenges for studying speciation in the marine environment. Our results build on a growing body of work that indicate the genus *Sebastes* is a good model for better understanding the mechanisms that promote speciation in the marine environment (Ingram 2011, Heras and Aguilar 2019, Behrans et al. 2021, Kolora et al. 2021). Using pooled exomes sequences, we were able to uncover islands of genomic differentiation in two different *Sebastes* species pairs that exhibit depth segregation. We were able to identify a greater number of islands in the *S. chlorostictus-S. rosenblatti* than the *S. crocotulus-S. miniatus* pair, which is expected given the disparity in divergence time observed in these taxa (Hyde and Vetter 2007). Enriched GO terms for genes found within the *S. chlorostictus-S. rosenblatti* species pair divergence islands suggest that genes involved in neuropeptide signaling and cellular homeostasis are important in the divergence of this species pair. A less clear pattern was observed for *S. crocotulus-S. miniatus*, but we were able to identify shared islands between the two species pairs. This work builds upon previous findings in the genus *Sebastes* and suggests a more complex pattern of genomic divergence as it relates to speciation in this group.

Exome wide analysis revealed a number of enriched GO terms found within significant outlier regions for the *S. chlorostictus-S. rosenblatti* comparison including cation homeostasis and the neuropeptide signaling pathway. The identification of genes in the neuropeptide signaling pathways in the *S. chlorostictus-S. rosenblatti* species pair is more directly related to ecological divergence and speciation in this group. Wang et al. (2021) point out the interplay between chemosensory divergence and ecological speciation. They focus on the importance of chemosensory drive in this process with a particular focus on diet adaptations. While dietary differences may exist in the *S. chlorostictus-S. rosenblatti* pair, it is likely adaptation to depth-related features are more important. Hyde and Vetter (2007) proposed a mechanism by which depth-segregation could lead to divergence in *Sebastes*. In *Sebastes*, juveniles are attracted to species-specific habitat types during settlement, and homing to a different depth-related habitat could contribute to sensory drive that would eventually lead to reproductive isolation via habitat differences (Heras et al. 2015).

We can only speculate on the relative importance of these pathways to the ecological divergence of this species pair. Genes involved in homeostatic functioning are likely crucial in maintaining cellular stability in the face of environmental differences. While there has not been adequate characterization of depth related habitat differences for any of the species we studied or the physiological demands of these environments, we can hypothesize that differences in temperature, pH, and salinity exist along the depth gradient that may drive local adaptation for these and other species. Overall, the enrichment of candidate genes near significant F_ST_ windows of genomic islands of differentiation for the *S. chlorostictus-S. rosenblatti* pair, provide functional descriptions of genes and gene networks related to behavior, development, homeostasis and immunity, supporting previous work on rockfish that has found similar evolutionary evidence corresponding to depth driving their speciation (Hyde and Vetter 2007, Ingram 2011, Burhens et al. 2021).

We found fewer enriched gene ontology groups for the *S. crocotulus-S. miniatus* species pair, concordant with the finding of fewer significant outlier windows. The significant gene ontology terms in this pair were associated with basic housekeeping functions (actin regulation of polymerization, the SWI/SNF pathways, histone lysine methylation pathways, tight junction and regulation of actin filament polymerization) and cannot be associated with differences in depth for this species pair. The SWI/SNF pathways act as ATP dependent chromatin remodelers that repress and activate genes and are associated with cardiovascular development (Supplementary Table S4). The finding that histone lysine methylation pathways are enriched in this species pair, could be related to hybrid sterility, as this pathway has been found to be related to hybrid sterility in mice (Mihola et al. 2009). Histone lysine methylation could indicate a postzygotic barrier in this species pair (Sha et al. 2020). It may be that the enriched genes found in this species pair are indicative of post zygotic barriers as chromatin remodeling genes are known to cause hybrid sterility and these molecular functions may be more present in longer diverged species of rockfish, however additional work needs to be done to assess the validity of these findings.

We found a limited number of shared outlier windows between the two species pairs, a window on chromosome 6 and one on chromosome 12. These overlapping windows contained only four annotated genes (Supplemental Table 4), some which are involved in basic metabolism. The lack of clear shared divergence islands across species pairs is likely due to the difference in estimated divergence times. This pattern was also observed by Behrens et al (2021) in a comparison of genome wide-divergence in three *Sebastes* species pairs and demonstrates the limitations of using F_ST_ in these types of studies (Quilodrán,et al. 2020).

### Comparison to Previous Work

A recent study of the genomic architecture of speciation in *Sebastes* found evidence for two ‘islands’ across two different chromosomes (Behrens et al. 2021). As in our study, Behrens et al. (2021) also utilized the *S. crocotulus-S. miniatus* pair and they found evidence for six regions of elevated genomic differentiation, however this study also utilized an additional species pair (*S. carnatus-S. chrysomelas*) that has a recent divergence (similar to *S. chlorostictus-S. rosenblatti)*. A more direct comparison of divergence islands with our study is not entirely possible, as Behrens et al (2021) used SNPs derived from reduced representation sequencing (ddRAD-Seq) and aligned their reads to a different genome assembly (*S. schlegelii*). The approaches to identifying the function of outliers was also different between the two studies, Beherens et al (2021) focused on identifying genes that contained outlier SNPs, while we looked at genes found within outlier windows and tested for the enrichment of GO terms for these genes. Regardless, we did not find similar gene sets in our analyses, with Behrens et al (2021) finding a set of genes related to vision and immune function. Nonetheless, two different methods to identify islands of differentiation show promise in identifying genomic mechanisms that drive the speciation of *Sebastes* and future work should focus on employing whole genome approaches and standardized genomic resources for *Sebastes* species.

### Demography of speciation

Examination of a limited set of demographic models indicated that the same model, secondary contact with asymmetric gene flow, was most likely for both species pairs. This suggests that the invasion of a novel habitat (deeper water in this case) is followed by a period of isolation in these species pairs. Models of ecological speciation predict that geneflow should persist upon invasion of the new ecological space. However, a recent study on depth segregated ecomorphs in *S. mentella* found support for demographic models that were similar to those found in this study (Benenstan et al. 2021). Benenstan et al. (2021) suggested that divergence in *S. mentella* was relatively recent (0.5 MYA) and driven by changes in sea level during the Pleistocene. Support for the secondary contact model found in this study is also supported by other studies of speciation history in marine organisms (Fairweather et al. 2018; Filatov et al. 2021; Leder et al. 2021). Another aspect of our analysis is that the directionality of asymmetric gene flow, following isolation, went consistently from the deeper species to the shallower species. It is unclear if this pattern will hold across other depth segregated species pairs in *Sebastes*, but could be indicative of climatic shifts impacting the depth distribution of these species. Future work on Northeastern Pacific *Sebastes* will determine if the overall pattern of isolation followed by secondary contact holds across depth segregated species pairs.

### Limitations

Our work is limited in that we utilized a pool-seq approach and only focused on enriched exome, CNE and UCE sequences. The main advantage of the pool-seq approach is that it reduces the overall cost of sequencing (Schlotterer et al. 2014). It does have the disadvantage that allele frequency estimates can be biased, but this is overcome with increased sequencing coverage (Schlötterer et al. 2016). We intentionally utilized high coverage regions in our SNP discovery steps (40-500 x coverage) to reduce any error. On top of this we utilized enriched sequences in our analysis, which has the advantage of reducing sequencing efforts to protein coding regions of the genome but would potentially be missing signals from extragenic regions.

### Conclusions

Our exome scan of two *Sebastes* species pairs revealed a handful of genes and pathways associated with depth-related divergence. There were a small number of shared islands of divergence between the pairs, but islands of divergence were more readily detected in the pair with more recent divergence. In the *S. chlorostictus-S. rosenblatti* pair we found enrichment for the neuropeptide synthesis pathway in outlier loci, which suggests that chemosensory drive may be involved in depth related speciation for this pair. Our analysis of demography of speciation revealed support for a similar model of divergence for the two pairs (isolation followed by secondary contact), which has been observed in other marine taxa. These results build on the growing knowledge of speciation history in the genus *Sebastes* and suggest *Sebastes* will continue to be a valuable model in understanding mechanisms of speciation in temperate marine fishes.

## ACKNOWLEDGMENTS

This project was supported by grants from the CSU Council on Ocean Affairs, Science & Technology (COAST-GDP-2019-001) and the National Science Foundation (DEB-2121145) to A. Aguilar. D. Olivares-Zambrano was supported by a CSU Council on Ocean Affairs, Science & Technology Graduate Student Research Award (CSUCOAST-OLIDAN-CSULA-AY1920) and by a grant from the National Institute of General Medical Sciences of the National Institute of Health (R25GM061331) (to K. Foster). Earlier drafts of this manuscript were improved with comments from K. Fisher and P. Krug (CSULA). We thank M. P. Harris at Harvard Medical School for technical laboratory support.

## DATA ACCESSIBILITY

All Illumina sequence data has been uploaded to the NCBI-SRA (project # PRJNA839756). All code for poolseq analysis is located on: https://github.com/aaguil67/aacode/blob/main/Sebastes_Olivares_1/Filtering_Mapping_Popoolation

