## Supplemental Information for "Speciation genomics and the role of depth in the divergence of rockfishes (*Sebastes*) revealed through Pool-seq analysis of enriched sequences"

Supplementary Table S1**:** Summary statistics for exome data

| Species | *S. chlorostictus* | *S. crocotulus* | *S. miniatus* | *S. rosenblatti* |
| --- | --- | --- | --- | --- |
| Raw Reads - Read 1 | 128,319,492 | 141,839,650 | 102,680,008 | 84,937,882 |
| Raw Reads - Read 2 | 128,319,492 | 141,839,650 | 102,680,008 | 84,937,882 |
| Total Raw Reads | 256,638,984 | 283,679,300 | 205,360,016 | 169,875,764 |
| Trim Reads - Read 1 | 124,311,746 | 137,520,573 | 99,147,354 | 82,548,830 |
| Trim Reads - Read 2 | 124,311,746 | 137,520,573 | 99,147,354 | 82,548,830 |
| Total Trim Reads | 248,623,492 | 275,041,146 | 198,294,708 | 165,097,660 |
| Properly Paired Reads Mapped to Reference Genome | 108,937,384 | 81,912,082 | 77,377,618 | 68,559,136 |
| Percentage of Properly Paired Reads Mapped to Reference Genome | 99.66% | 99.12% | 99.35% | 99.73% |
| Percentage of Average Coverage Mapped to Reference Genome | 35.26% | 30.94% | 29.53% | 28.64% |
| Percentage of Average Depth Mapped to Reference Genome | 12.07% | 10.04% | 9.90% | 9.60% |

Supplementary Table S2: Significant windows of differentiation for the *S. chlorostictus* and *S. rosenblatti* comparison. Highlighted windows are shared with the *S. crocotulus-S. miniatus* comparison.

| Chromosome # | NCBI Accession # | Start Bp | End Bp |
| --- | --- | --- | --- |
| 1 | NC_051269.1 | 5225191 | 5356807 |
| 1 | NC_051269.1 | 35467223 | 36336323 |
| 1 | NC_051269.1 | 41553085 | 41875484 |
| 1 | NC_051269.1 | 42060235 | 42115804 |
| 1 | NC_051269.1 | 42578894 | 43876219 |
| 2 | NC_051270.1 | 212739 | 1023943 |
| 3 | NC_051271.1 | 11249438 | 11273040 |
| 5 | NC_051273.1 | 752477 | 1347961 |
| 5 | NC_051273.1 | 35105744 | 35693947 |
| 6 | NC_051274.1 | 28125426 | 28146180 |
| 6 | NC_051274.1 | 28366997 | 28863953 |
| 6 | NC_051274.1 | 33908745 | 33922407 |
| 6 | NC_051274.1 | 33922479 | 33932037 |
| 10 | NC_051278.1 | 4780440 | 4780556 |
| 11 | NC_051279.1 | 14331294 | 14381322 |
| 12 | NC_051280.1 | 12165513 | 12521340 |
| 12 | NC_051280.1 | 14666841 | 14716987 |
| 12 | NC_051280.1 | 32492987 | 32497311 |
| 13 | NC_051281.1 | 4230382 | 4230862 |
| 13 | NC_051281.1 | 6573222 | 6625313 |
| 14 | NC_051282.1 | 29798867 | 30009813 |
| 16 | NC_051284.1 | 1041715 | 1462337 |
| 17 | NC_051285.1 | 18110333 | 18918230 |
| 18 | NC_051286.1 | 4233137 | 4305852 |
| 18 | NC_051286.1 | 26386754 | 27537508 |
| 18 | NC_051286.1 | 28337654 | 29095057 |
| 20 | NC_051288.1 | 13091065 | 13185450 |
| 20 | NC_051288.1 | 24378485 | 24430264 |
| 21 | NC_051289.1 | 4458558 | 5180472 |
| 21 | NC_051289.1 | 7299259 | 7301243 |
| 22 | NC_051290.1 | 11184406 | 11215966 |
| 22 | NC_051290.1 | 17042459 | 17358868 |
| 23 | NC_05129.1 | 9484382 | 9886423 |

Supplementary Table S3: Significant windows of differentiation for the *S. crocotulus* and *S. miniatus* comparison. Highlighted windows are shared with the *S. chlorostictus* and *S. rosenblatti* comparison.

| Chromosome # | NCBI Accession # | Start Bp | End Bp |
| --- | --- | --- | --- |
| 6 | NC_051274.1 | 28425422 | 28930966 |
| 8 | NC_051276.1 | 34451820 | 35437058 |
| 11 | NC_051279.1 | 15036147 | 15877357 |
| 12 | NC_051280.1 | 4214238 | 4463848 |
| 12 | NC_051280.1 | 31843674 | 32493378 |
| 14 | NC_051282.1 | 17769519 | 17967545 |
| 14 | NC_051282.1 | 24614709 | 25366241 |
| 15 | NC_051283.1 | 8527532 | 9385230 |
| 18 | NC_051286.1 | 2714834 | 3035398 |
| 21 | NC_051289.1 | 23840005 | 24582084 |

Supplemental Table S4: Shared genes found in overlapping Fst outlier windows across the two species pairs

| Gene ID | Gene Name | Function |
| --- | --- | --- |
| si:dkey-96f10.1 | 6-phosphofructo-2-kinase/fructose-2 2C6-bisphosphatase 2C | glycolysis |
| znf622 | zinc finger protein 622 | zinc ion binding activity |
| myo10 | myosin X | ATP binding activity; actin binding activity |
| adcy8 | adenylate cyclase 8 (brain) | axon midline choice point recognition and retinal ganglion cell axon guidance |

Supplementary Table S5: Bonferroni corrected candidate genes and their functions by species pair.

| Species Pair | Candidate Gene | Gene Name | Function | Link | Reference |
| --- | --- | --- | --- | --- | --- |
| S. chlorostictus-S. rosenblatti | adcyap1b | Adenylate cyclase activating polypeptide 1b | Activates G protein and stimulates pituatary cell release of adenylate cyclase. Neuron projection. Stimulates melophore production | https://www.uniprot.org/uniprot/Q9NUT2 | The Uniport Team, 2021 |
|  | abcb8 | ATP binding cassette subfamily B member 8 | Binds ATP, as a subunit of the mitochondrial potassium channel and is located in the innter mebrane. Mediates potassiun currents along the inner mebrane and is required for cardiac maintence. Influences iron trasnport in the mitochondria and is required. | https://www.ncbi.nlm.nih.gov/gene/514 | Sayers et al. 2022 |
|  | atp5f1e | ATP synthase F1 subunit epsilon | Mitochondrial ATP synthase that catalizes ATP synthesis/ | https://www.ncbi.nlm.nih.gov/gene/51606 | Sayers et al. 2022 |
|  | atp6v1h | ATPase H+ Transporting V1 Subunit H | Mediates accidification in intracellular organelles. | https://zfin.org/ZDB-GENE-010718-1#summary | Ruzicka et al. 2019 |
|  | atp1b2b | ATPase Na+/K+ transporting subunit beta 2b | Contributes to sodium:potassium exhange ATPASE activity. Involved in development of heat, sensory organs and startle response. | http://zfin.org/ZDB-GENE-090312-136#summary | Ruzicka et al. 2019 |
|  | atp6ap1a | ATPase H+ transporting accessory protein 1a | proton transport atpase activity. Regulation of pH. Human orthologues implicated in primary immunodeficiency disease. | https://zfin.org/ZDB-GENE-040426-2222 | Ruzicka et al. 2019 |
|  | ca15b | Carbonic anhydrase XVb | Carbonate dehydratase activity and zinc ion binding. Germ cell migration and intracellular pH regulation. | https://zfin.org/ZDB-GENE-140624-1 | Ruzicka et al. 2019 |
|  | galr1b | Galanin receptor 1b | Galanin receptor activity and involved in G-protein coupled receptor activity | https://medlineplus.gov/genetics/gene/itgb4/ | MedlinePlus, 2020 |
|  | itb4r | Integrin beta subunit | A receptor for laminin. | https://www.uniprot.org/uniprot/P16144#function | The Uniport Consortium, 2021 |
|  | itb4r2a | Integrin beta subunit | A receptor for laminin. | [https://www.uniprot.org/uniprot/P41145https://](https://www.uniprot.org/uniprot/P41145https:/) | The Uniport Consortium, 2021 |
|  | oprk1 | Opiod receptor kappa 1 | Mediates stress responses. Pain perception. Arousal and regulation of autonomic nervous system and neuroendorcrine functions. | https://zfin.org/ZDB-GENE-040114-1 | Ruzicka et al. 2019 |
|  | pnocb | Prepronociceptin b | Involved in ectodermal placode development. | https://www.uniprot.org/uniprot/Q96A98 | The Uniport Consortium, 2021 |
|  | pth2 | Parathyroid horome 2 | May inhibit cell proliferation. May have a role in spermatogenesis as a neuropeptide. | https://pubmed.ncbi.nlm.nih.gov/33044947/ | Sayers et al. 2022 |
|  | slc4a2b | Solute carrier family 4 member 2b | Involved in hair cell development. | http://www.informatics.jax.org/marker/MGI:1933532 | Bult et al. 2019 |
|  | slc12a9 | Solute Carrier Family 12 Member 9 | potassium chloride transport | https://www.genecards.org/cgi-bin/carddisp.pl?gene=SLC2A10 | Safran et al. 2022 |
|  | slc2a10 | Solute carrier family 2 member 10 | Glucase transport. Required for development of cardiovascular system | https://www.genecards.org/cgi-bin/carddisp.pl?gene=SLC9A7 | Safran et al. 2022 |
|  | slc9a7 | Solute Carrier Family 9 Member A7 | Mediates exchange of potassium and calcium in endosomes. Regulation of volume and pH in golgi apparatus | https://zfin.org/ZDB-GENE-130531-48 | Ruzicka et al. 2019 |
|  | tap2t | Transporter associated with antigen processing, subunit type t, teleost specific | Contribute to MHC class I protein binding activity | https://zfin.org/ZDB-GENE-030131-5725 | Ruzicka et al. 2019 |
| S. crocotulus-S. miniatus | arid1ab | AT rich interactive domain 1Ab (SWI-like) | Contributes to nucleosome binding acitivity/ ATP-dependent chromatin remodeling. | https://www.uniprot.org/uniprot/Q8NFD5#function | The Uniport Consortium, 2021 |
|  | arid1b | AT-rich interaction domain 1B | belongs to neuroprogenator specific chromatin remodeling complex (Nbaf) | https://www.uniprot.org/uniprot/Q9UQB8 | The Uniport Consortium, 2021 |
|  | baiap2a | Brain-specific angiogenesis inhibitor 1-associated protein 2 | Cytoskeleton anchor activity. Neurite growth. Reorganization of actin cytoskeleton in response to bacterial infection. Filopodia formation. | https://zfin.org/ZDB-GENE-040704-22 | Ruzicka et al. 2019 |
|  | cdc42ep4a | CDC42 effector protein (Rho GTPase binding) 4a | GTP-Rho binding activity. | https://www.uniprot.org/uniprot/Q6DBT9 | The Uniport Consortium, 2021 |
|  | cldnk | Claudin k | Calcium independent cell adhesion activity that leads to tight-junction specific removal of the intracellular space. | https://www.ncbi.nlm.nih.gov/gene/1786 | Sayers et al. 2022 |
|  | dnmt1 | DNA methyltransferase 1 | Encodes enzyme that transfers methyl groups to cytosine. | https://www.uniprot.org/uniprot/P51153 | The Uniport Consortium, 2021 |
|  | rab13 | RAB13, member RAS oncogene family | Key regulators of intracellular membrane trafficking. More broadly play a role in the establishment of setoli cellbrarrier, angiogenesis, neurite outgrowth and glucose homeostasis. | https://www.uniprot.org/uniprot/O15047 | The Uniport Consortium, 2021 |
|  | setd1a | SET Domain Containing 1A, Histone Lysine Methyltransferase | Catalyzes methyl group transfer to 'LS-4' histone H3. Broadly involved in transcriptional programming of inner mass stem cells and neuroprogenators. | https://www.uniprot.org/uniprot/Q08BR4 | The Uniport Consortium, 2021 |
|  | setdb1b | Histone-lysine N-methyltransferase SETDB1-B | Trimethylates 'LS-9' of histone H3. Central role in silencing of eurochromatin genes. Hypermethylation and silencing of tumor suppresor genes. Maintains trancriptionally repressive state of genes in undifferentiatied embryonic stem cells. | https://www.uniprot.org/uniprot/Q9BTW9 | The Uniport Consortium, 2021 |
|  | tbcd | Tubulin Folding Cofactor D | Regulation of microtubule polymerization or depolymorization. Involved in neuron morphogenesis. | https://www.uniprot.org/uniprot/P0C1Z6#function | The Uniport Consortium, 2021 |
|  | tfpt | TCF3 fusion partner | Promotes apoptosis. Regulatory component of the chromotin remodeling INO80 Complex | https://www.uniprot.org/uniprot/P0C1Z6#function | The Uniport Consortium, 2021 |
|  | zgc:158689 | Uncharacterized LOC117533444 | Involved in actin-filament organization. Positive regulation of cytoskeletan organization | https://zfin.org/ZDB-GENE-070112-1902 | Ruzicka et al. 2019 |
